# In Situ Biocatalytic Color Generation Using Living Whole-Cell Systems for Sustainable Textile Dyeing

**DOI:** 10.1101/2025.11.02.684288

**Authors:** Asish Das, Tanu Tiwari, Karthik Pushpavanam

## Abstract

Pigments are widely utilized in industries such as textiles, cosmetics, food, and packaging. However, conventional synthetic pigment production relies on petroleum-based feedstocks, consumes significant energy, and involves toxic chemicals raising environmental and safety concerns. Natural pigments offer a sustainable and biodegradable alternative, yet their large-scale application is hindered by limited availability, batch variability, and geographical dependence. To overcome this, microbial biosynthesis has attracted attention as a controllable and scalable route for pigment production independent of environmental fluctuations. Nevertheless, most existing approaches retain multistep workflows – pigment synthesis, extraction, purification, and application. This compromises process efficiency and reproducibility. Herein, we present a whole-cell biocatalytic approach for melanin synthesis and deposition, wherein recombinant E. coli expressing tyrosinase is employed to convert L-tyrosine into melanin through copper-dependent oxidative polymerization. By harnessing intact microbial cells as self-contained biocatalytic units, this system bypasses the need for enzyme extraction and purification. Prior to applying the culture media containing melanin on material surfaces such as cotton and wood, the process conditions were optimized to enhance melanin yield. Subsequently, the material surfaces incubated in the melanin culture medium were characterized for their surface morphology and chemical modifications through scanning electron microscopy, Fourier transform infrared spectroscopy, and X-ray photoelectron spectroscopy. The color fastness properties of the material were also evaluated in the presence of water and detergents and subsequently improved through post-treatment processes. In addition, the melanin-coated cotton demonstrated enhanced photothermal performance compared to uncoated controls. All these taken together, this work establishes a simplified and potentially scalable route toward sustainable pigment production and direct application through enzyme-driven in situ biocatalysis on various material surfaces.

## INTRODUCTION

Pigments are essential functional components employed across diverse sectors including textiles, cosmetics, food, and packaging.^1^ They represent a market valued at USD 38.2 billion in 2022 and projected to grow at over 5% annually by 2030.^1–3^ These compounds include structurally and functionally distinct chemical classes including but not limited to carotenoids, tetrapyrroles, and flavonoids.^4–7^ However, industrial synthesis is energy intensive, dependent on petroleum-derived feedstocks and involves toxic reagents raising sustainability concerns.^8^ These limitations collectively highlight the urgent need for alternative, environmentally responsible approaches to pigment production and application.

Natural pigments, derived from plants, insects, animals, and minerals, span a color palette from yellow to black with light absorption in the visible range (400 to 800 nm).^9–11^ Historically significant examples include madder red, blue indigo, saffron or turmeric yellow, alongside mineral pigments such as ochre, limestone, and charcoal.^12–17^ Recent technological advances have improved their accessibility. For instance, Noman Habib et al. recovered colorants from *Curcuma aromatica* using a microwave-assisted extraction method and applied them to silk resulting in enhanced color depth and fastness.^18^ Luis Eduardo Ordóñez-Santos optimized the ultrasound-assisted extraction of carotenoids from mandarin epicarp and demonstrated their application as natural colorants.^19^ Daniel López-Rodríguez investigated the extraction of natural pigments from various plant materials using different extraction methods such as freezing-assisted aqueous extraction, heat-assisted ethanol extraction, and boiling water extraction and characterized their properties for potential application as natural colorants.^20^ Muhammad Ibrahim et al. extracted the dye from Bixa orellana (annatto) seeds, using microwave-assisted extraction for silk dyeing with added functional properties.^21^ Despite these advances, natural pigment production remains constrained by seasonal variability, geographical dependence, and limited scalability, posing challenges for industrial implementation.^22^

Microbial biosynthesis has emerged as a promising platform to address these constraints, offering scalability, process tunability and independence from environmental fluctuations.^23^ Notably, microorganisms can be engineered to synthesize pigments characteristic of plant and animal sources.^24^ Yajun Yan et al engineered a four-enzyme plant biosynthetic pathway in *E*.*coli* to convert flavanones into glycosylated anthocyanins.^25^Stephen Hart et al. engineered *E*.*coli* with a cloned *Rhodococcus* sp. fragment to produce indigo and indirubin from tryptophan-derived indole.^26^ At the industrial scale, Colorifix, has demonstrated biologically mediated textile coloration with reduced environmental impact.^27^ Nevertheless, most existing approaches retain multistep workflows – pigment synthesis, extraction, purification and application.^28^ This compromises process efficiency and reproducibility.

Among naturally occurring pigments, melanin is particularly attractive for functional applications.^29^ This biopolymer is formed through the oxidative polymerization of phenolic or indolic compounds and is widely distributed across microorganisms, plants, and animals.^30^ Its widespread appeal is due to its unique multifunctional properties, including broadband light absorption, photothermal conversion, radical scavenging, and strong metal-ion binding.^31–33^ Soo-Yeon Ahn et al. produced melanin using *Streptomyces glaucescens* and engineered *E*.*coli and* applied it to cotton with improved color fastness.^34^ Esfandiar Pakdel et al. extracted melanin from yak hair and applied it to cotton to develop a multifunctional coating with superhydrophobicity and UV protection.^35^ Wenjing Liu et al. synthesized silver-loaded melanin-like poly(levodopa) nanoparticles via a one-step method and applied them to cotton to impart UV protection, photothermal activity, and antibacterial functionality while maintaining breathability.^36^ However, these approaches share common drawbacks including the requirement and reliance on pre-synthesized or externally processed pigments.^37^ In this context, strategies enabling *in situ* pigment generation directly on the target material surface offer a compelling alternative.

Herein, we present a biocatalytic approach for melanin synthesis and deposition, wherein tyrosinase-catalyzed oxidation of L-tyrosine in the presence of copper ions generates reactive intermediates that undergo polymerization to form melanin. This system enables simultaneous pigment formation and material surface coating, thereby eliminating the need for pigment extraction and transfer steps. The resulting coatings are directly formed on cotton and wood under optimized conditions, exhibiting uniform deposition and photothermal performance. This work establishes a simplified, sustainable and potentially scalable route through whole cell mediated biocatalytic production and in situ application on the material surface.

## MATERIALS AND METHODS

### Materials

Cotton and detergent were outsourced from local market. Chir wood (Pinus roxburghii) was acquired from the mechanical workshop at IIT Gandhinagar. We sourced the materials from the following suppliers. **SRL Pvt. Ltd**. Methanol, Cupric Sulfate Pentahydrate, and Luria Bertani (LB) broth. **Tokyo Chemical Industry (TCI) Pvt. Ltd**. L-tyrosine. **Hi Media Laboratories**. Isopropyl-ß-D-1-thiogalactopyranoside (IPTG) and Kanamycin sulfate. **Addgene**. Plasmid pLN-AIDA-Tyr1 (Plasmid #192831) **(Table S1). Supelco**. Isopropyl alcohol (IPA), Ethanol. **Sigma Aldrich**. Sodium carbonate, Tannic acid, Potassium aluminum sulfate (potash alum), Acetic acid, and Sodium Hydrosulfite. **Loba**. Sodium Hydroxide. **Actylis Lab Solution**. Hydrogen Peroxide.

### Expression and characterization of the melanin

The pLN-AIDA-Tyr1 obtained as agar stab was transformed into *Escherichia coli* BL21(DE3) cells. The transformed colonies were isolated in LB media supplemented with kanamycin (50 mg/mL). 1% inoculum of the overgrown culture of *Escherichia coli* BL21(DE3) was used to inoculate 100 mL culture with kanamycin. The culture was incubated at 37 °C with continuous shaking until it reached an OD_600_ between 0.8 - 1. Melanin production was induced by adding 1 mM IPTG, 0.6 mM tyrosine and 30 µM CuSO_4_·5H_2_O. The culture was further grown at 37 °C for 24 h. The cells were harvested by centrifugation (6000 RPM, 15 min, 4°C) and the supernatant and pellets were stored at 4 °C and -20 °C respectively until further use.

### Coating of cotton

The cotton samples were cut into pieces of dimensions 2 × 2 cm. The samples were pretreated by first dipping them into a 16.7 mM acetic acid solution at 85 °C for 20 minutes. Subsequently, the samples were dipped in a solution containing 25 mM NaOH and 30 mM H_2_O_2_ at 95 °C for 60 min, then rinsed thoroughly in 16.7 mM acetic acid at 50 °C for 10 min. After that, it was dipped into a solution containing 42 mM Na_2_S_2_O_4_ and 56 mM Na_2_CO_3_ at 80 °C for 30 minutes. These steps were performed to ensure that the cotton is free of any chemicals, waxes or oils and ready for further treatment. Then, the cotton samples were treated with 10% weight of fabric tannic acid at 80 °C for 1 h, followed by treatment with 15 % potash alum for 1 h. Finally, the cotton was immersed in the melanin culture and heated at 100 °C for 45 mins. Finally, acetic acid was added to the same mixture and kept for 15 min to coat the cotton.

The samples were then heat-pressed on a flat plate at 80 °C, after which they were thoroughly washed with water. An extra coating of melanin was then applied on the samples by soaking them in the melanin culture media and heating them up at 100 °C for 45 min. This coating process was repeated for three cycles.

### Coating of wood

The chir wood was cut into small blocks with dimensions 1 × 1 × 1 cm. The wood was sterilized by rinsing it with 100% IPA, followed by drying at 100 °C for 15 min to remove the residual moisture. The wood was also sterilized in UV for 15 min before every experiment. The sterilized wood pieces were added to the culture during the induction stage after adding the IPTG, tyrosine and CuSO_4_·5H_2_O and kept at incubation at 37 °C while shaking.

### Scanning electron microscopy

The uncoated and coated cotton samples were dried for 24 hours at 60 □. Also, for the uncoated and melanin-coated wood, the samples were scraped and washed with water to remove any residual components on the surface. These wood samples were dried at 60 □ for 12 h to remove the excess moisture. Platinum was used to coat the dried cotton samples for 60 seconds, while gold was used for conducting the dried wood samples for 180 seconds. The micrographs were recorded under a field-emission scanning electron microscope (FE-SEM) of JEOL JSM-7600F in SEI mode.

### Fourier Transform Infrared Spectroscopy

The cotton samples were prepared by drying them at 60 □ for 24h. The wood samples were prepared by scraping the wood into a dimension of 1 cm× 1cm× 1mm. The spectra were recorded using a Perkin Elmer FTIR spectrophotometer across the range of 4000−500 cm^-1^.

### X-ray Diffraction of wood

The wood samples were prepared by scraping the wood in the dimensions of (1 cm× 1cm× 1mm). XRD measurements were done using the Rigaku SmartLab 9KW, operated with Cu Kα radiation (λ□=□1.5418□Å) at a scan rate (2θ) of 5° min^−1^. The incident angle was 1° (⍰ = 0.5°) with a step size of 0.05, ranging from 10° to 60°. The melanin coated wood and uncoated wood (control) were analyzed under XRD, maintaining a thickness of 1 mm.

### X-ray Photoelectron Spectroscopy of uncoated and coated material surfaces

The cotton samples were prepared by drying them at 60 □ for 24 h. The wood was scraped out to a dimension of 1 cm× 1 cm× 1 mm and dried at 60 °C for 12 h. The XPS spectra were recorded for both samples by an X-ray photoelectron spectrometer (K-Alpha). The X-ray source used was Al-Kα micro focused monochromator with variable spot size (50-400) µm. The peaks for all the elements were deconvoluted using the Origin Pro and Avantage software.

### Mechanical characterization of wood by UTM

The melanin-coated wood was dried at 100 °C for 12 h. The compression test was performed with the help of a Universal Testing Machine (UTM) Model No. CMT5205. The samples were mounted and subjected to a displacement rate of 5 mm min^-1^ until a displacement of 5 mm was achieved. Young’s modulus was determined by calculating the slope (Change in Stress / Change in Strain) of the linear elastic region of the stress-strain curve.

### Evaluation of the photothermal performance

The photothermal performance of both melanin-coated cotton and wood and respective controls (without melanin coating) was evaluated. All the samples were irradiated using a laser with a wavelength of 808 nm. Each sample was exposed to the laser for defined time intervals, with the beam directed onto the surface. The resulting temperature changes were recorded using an infrared thermal camera (FLIR E5).

### Statistical analysis

All experiments were performed in three independent samples, and data are presented as mean ± standard deviation (SD).

## RESULTS AND DISCUSSION

Cotton is a natural, polymer-based textile composed of cellulose fibers rich in surface hydroxyl groups, which impart inherent hydrophilicity to the fabric.^38,39^ Owing to its wide availability, low cost, biodegradability, softness, and comfort, it remains one of the most extensively used textile materials worldwide accounting for over 82% of global natural fiber consumption.^40,41^ Traditionally, cotton coloration has predominantly depended on synthetic dyes and pigments because of their broad color spectrum, ease of processing, and economic feasibility.^42^ However, growing environmental concerns surrounding synthetic colorants have necessitated the development of sustainable alternatives for textile coloration. In this context, we have developed a whole-cell-mediated biocatalytic pigment production process for the simultaneous coloration of cotton during pigment synthesis.

To implement the whole-cell biocatalytic strategy described above, the plasmid construct pLN–AIDA–Tyr1 was engineered for tyrosinase expression **(Figure 1A)**. Melanin synthesis is induced by isopropyl β-D-1-thiogalactopyranoside (IPTG, 1 mM), with tyrosine serving as the substrate and CuSO_4_ acting as the cofactor required for tyrosinase activity and subsequent melanin production (**Figure 1B**). The production of tyrosinase by the tyr gene was confirmed by the visible color change of the culture, which turned black in contrast to the yellow appearance of the growth media **(Figure 1C)**. The reaction schematic illustrates the melanin biosynthetic pathway initiated from tyrosine, progressing through the intermediates L-DOPA, dopaquinone, and dopachrome **(Figure 1D)**.^37,43^ After induction, melanin accumulation is readily visible in both the cell pellet and the supernatant (**Figure 1E**). SDS-PAGE revealed a protein band around 37 kDa corresponding to tyrosinase, confirming the expression of the enzyme in the culture **(Figure 1F)**.

**Figure 1.**
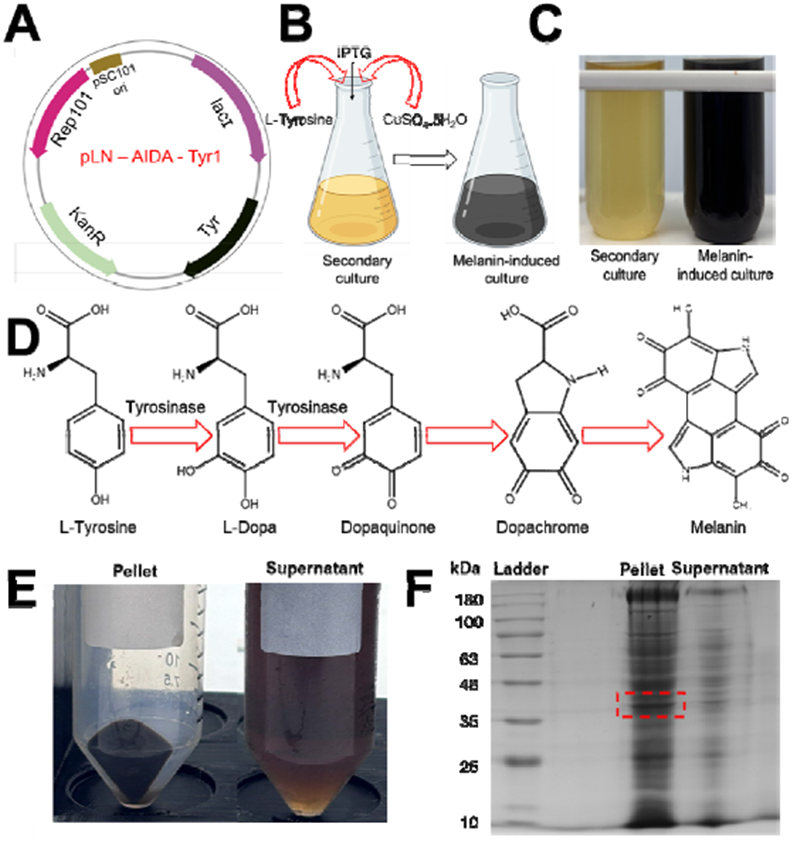
Expression of tyrosinase and production of melanin. **(A)** Schematic representation of plasmid construct pLN-AIDA-Tyr1, **(B)** Schematic representation of IPTG induction for melanin production in the presence of tyrosine as a substrate and CuSO_4_.5H_2_O as a cofactor, **(C)** Digital image of the LB media taken prior to melanin production with yellow coloration (left) and the image taken after the IPTG induction with melanin production with black coloration (right), **(D)** The biosynthetic pathway for the conversion of tyrosine into melanin, **(E)** The melanin produced is centrifuged and the corresponding pellet and the supernatant appears black in color and **(F)** SDS-PAGE analysis of expressed melanin including lane 1: molecular weight marker (kDa), lane 2: pellet of expressed melanin culture and lane 3: supernatant of expressed melanin culture. The highlighted white dashed box indicates the tyrosinase enzyme around 37 kDa.

We assessed the growth profile of the bacteria by tracking the optical density (OD_600_) over time prior to subsequent application (**Figure 2A**). IPTG induction was performed when the OD_600_ reached the target optimal range of 0.6–0.8, corresponding to the mid-exponential growth phase, to ensure maximum tyrosinase expression and melanin production. The expression of tyrosinase in the presence and absence of IPTG was investigated using SDS-PAGE analysis **(Figure 2B)**. The gel exhibited a prominent band at the expected molecula weight (37 kDa) in the protein pellet in the presence of IPTG, whereas in the absence of IPTG, weaker expression along with additional non-specific protein bands was observed. Additionally, a faint band corresponding to approximately 37 kDa was also detected in the respective supernatants.

**Figure 2.**
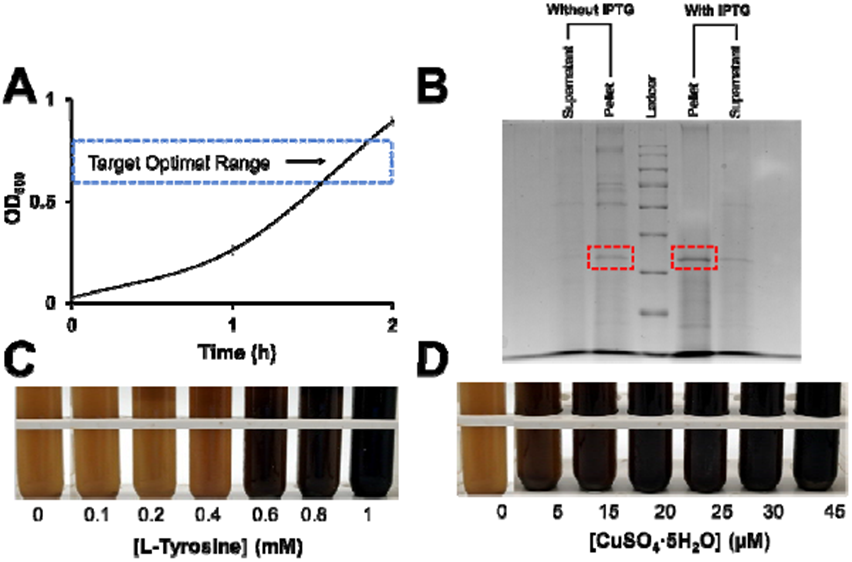
Optimization and analysis of melanin production. **(A)** Growth kinetics of recombinant *E. coli* monitored by OD□□□ measurements over time, **(B)** SDS–PAGE of the expressed tyrosinase, where lane 2 and lane 3 represent the supernatant and corresponding pellet of the culture medium without IPTG induction, respectively, lane 4 corresponds to the protein molecular weight marker, and lane 5 and lane 6 represent the pellet and corresponding supernatant of the culture medium with IPTG induction, respectively. The highlighted regions indicate the expected band of tyrosinase (37 kDa), **(C)** Digital images of the culture media following melanin production at constant IPTG (1 mM) and CuSO_4_·5H_5_O (30 µM) concentrations, while varying the concentration of L-tyrosine from 0 to 1 mM. and **(D)** Digital images of the growth media following melanin production at constant IPTG (1 mM) and L-tyrosine (0.6 mM) concentrations, while varying the concentration of CuSO_4_·5H_2_O from 0 to 45 µM.

We also investigated the role of tyrosine and CuSO_4·_5H_2_O during melanin production. The black coloration due to melanin was optimized, varying the concentrations of L-tyrosine and CuSO_4_. L-tyrosine acts as the substrate in the tyrosinase-catalyzed pathway, where tyrosinase facilitates the oxidation of L-tyrosine to L-DOPA in the presence of oxygen, followed by further oxidation to dopachrome, which subsequently undergoes spontaneous polymerization to form melanin.^44,45^ Cuprous ions act as a cofactor for tyrosinase, facilitating the conversion of dopachrome into 5,6-dihydroxyindole-2-carboxylic acid (DHICA).^46,47^ We varied the tyrosine concentration from 0.1 mM to 1 mM while IPTG (1 mM) and CuSO_4_5H_2_O (30 µM) were held constant. We observed a clear change in color in the media from yellow to black with an increasing concentration of tyrosine. The culture displayed a brown coloration at tyrosine concentrations up to 0.4 mM, while higher concentrations resulted in the formation of black pigmentation, with the intensity of melanin increasing proportionally with increasing tyrosine concentration **(Figure 2C, S1A and S1B)**. A similar trend was observed upon varying the concentration of CuSO_4_·5H_2_O from 5 µM to 45 µM while maintaining constant concentrations of IPTG (1 mM) and tyrosine (0.6 mM), indicating enhanced melanin production with increasing copper concentration. **(Figure 2D, S1C and S1D)**. The media was dark brown till the concentration of 5 µM, and then it turned black with increased intensity. Based on these observations, the subsequent experiments were performed using optimized concentrations of IPTG (1 mM), tyrosine (0.6 mM), and CuSO_4_·5H_2_O (30 µM).

The synthesized melanin was characterized through UV–Visible spectra (200–700 nm) which demonstrated the characteristic broad-band monotonic absorbance profile for both the pellet and the corresponding supernatant **(Figure S2A)**. The broadband absorption behavior arises from the intrinsic molecular structure within melanin aggregates.^48^ Subsequently, the FTIR spectra for the lyophilized pellet was determined to identify the functional groups **(Figure S2B)**.^49^ The FTIR spectra exhibited six characteristic peaks, with a broad peak centered at 3382 cm^-1^ corresponding to the O–H and N–H stretching vibrations. The peak observed at 2931 cm^-1^ is attributed to aliphatic C–H stretching vibrations. Furthermore, the peaks at 1636 cm^-1^ and 1533 cm^-1^ corresponded to aromatic C=C stretching and N–O stretching, respectively.

Subsequently, the bleached cotton was heated in the melanin culture medium at ∼100 °C, resulting in its black coloration **(Figure 3A)**. However, after 24 h drying, noticeable fading in color intensity was observed compared to the freshly treated samples. The observed fading is likely due to the diminished affinity between melanin and cotton.^50^ To improve the melanin retention, we pre-treated the cotton samples with tannic acid, potash alum and acetic acid. This treatment resulted in an intense black coating on the cotton, which remained clearly visible after drying. We posit that the pre-treatment resulted in improved affinity for the uptake of the melanin due to the introduction of additional sites over its surface. Tannic acid and potash alum acts as mordants, promoting stronger interactions with the cellulose and thereby enhancing melanin uptake.^51^ Tannic acid adheres to the cellulose via hydrogen bonds creating a polyphenol-based primary layer with π-π interactions with the aromatic structures of melanin.^52,53^ This layer acts as a molecular bridge that promotes the interaction of melanin onto the cotton surface. Simultaneously, potash alum facilitates metal–ion coordination interactions between tannic acid, melanin, and Al^3^□ ions, further enhancing melanin fixation and color uptake.^54^ Additional control experiments were performed to indicate that all three ingredients are required for successful coating **(Figure S3)**.

**Figure 3.**
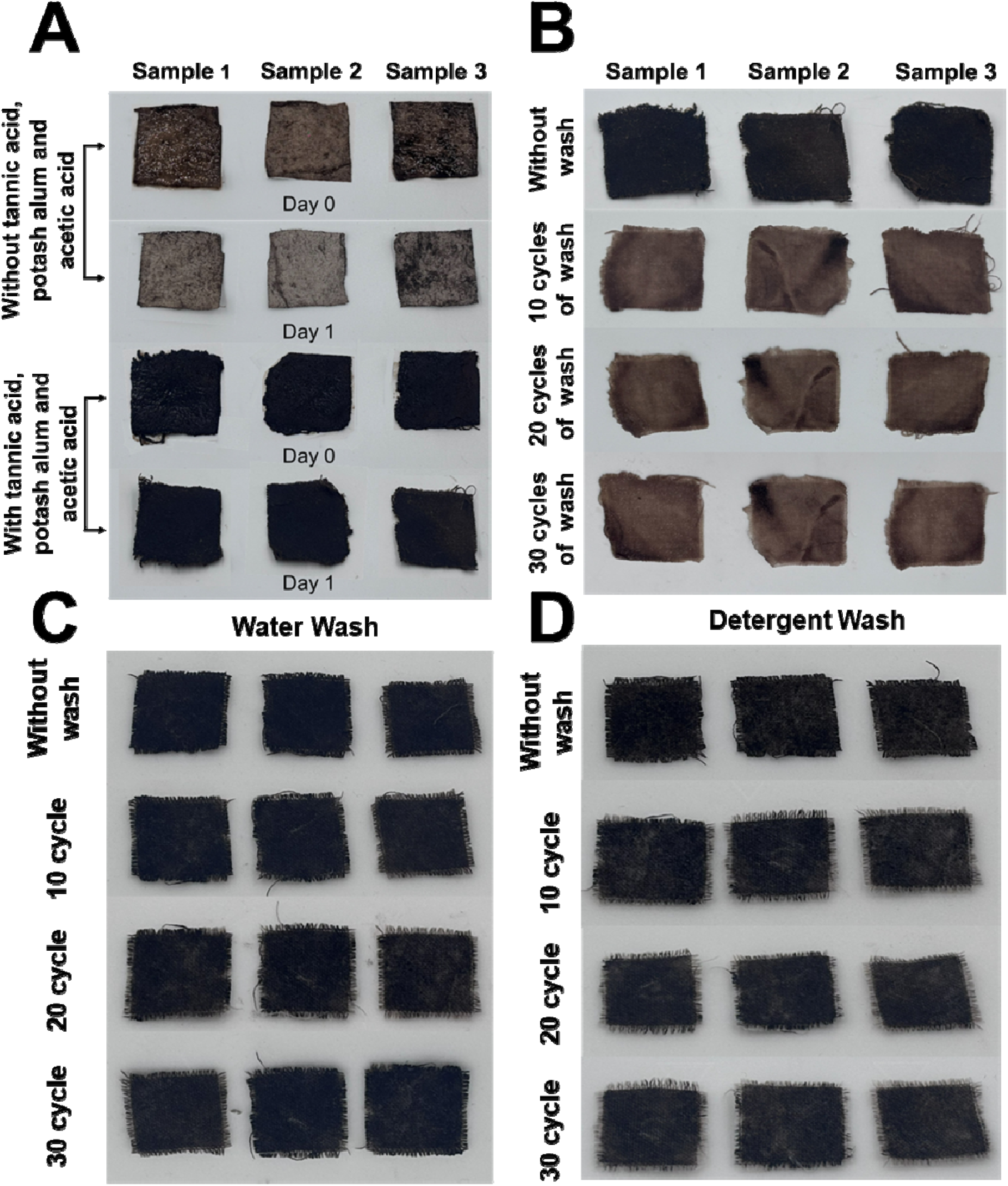
Melanin-mediated coating and color fastness of cotton. **(A)** Digital images showing the coating behavior of cotton treated with melanin culture in the absence and presence of mordant pre-treatment. The upper row represents cotton samples coated without pre-treatment, exhibiting noticeable fading after drying for 24 h compared to the freshly treated samples. The lower row represents cotton samples pre-treated with tannic acid, potash alum, and acetic acid prior to melanin coating, displaying intense and stable black coating even after drying under identical conditions, **(B)** Digital images demonstrating the color fastness of melanin-coated cotton samples pre-treated with tannic acid, potash alum, and acetic acid. The top row represents the freshly coated cotton samples prior to washing, while the subsequent rows correspond to samples subjected to 10, 20, and 30 water-washing cycles, respectively. A clear decrease in color intensity was observed with increasing washing cycles, **(C)** The post-treated coated cotton samples retained their black coating after 30 washing cycles with water and **(D)** The coated cotton samples also exhibited strong resistance to color after 30 washing cycles in the presence of detergent, demonstrating improved color fastness.

The color fastness of melanin on the cotton surface was evaluated through repeated water-washing cycles. After 30 washing cycles, a reduction in the intensity of the black coating was observed **(Figure 3B)**. Additionally, a noticeable color loss was observed after the first wash cycle with detergents indicating poor washing stability of the coated melanin **(Figure S4)**. To overcome this, a post-treatment including three successive layers of melanin coating with heat pressing after each melanin coating cycle was employed. We hypothesized that the heat press process could result in partial loosening and swelling of the cellulose network, facilitating greater penetration and absorption of melanin into the structures.^55,56^ As anticipated, the pre-treatment facilitated enhanced stability for 30 cycles of washing using water and detergent containing solutions **(Figure 3C and 3D)**.

The surface morphology of the uncoated cotton along with the melanin-coated cotton was determined by scanning electron microscopy. The uncoated cotton depicted smooth, well-defined cylindrical structures arranged in a tightly bundled architecture (**Figure 4A and S5A)**.^57^ In contrast, the SEM image of the melanin-coated cotton revealed similar cylindrical structures with a non-homogenous surface likely due to melanin coating **(Figure 4B and S5B)**. Subsequently, FTIR spectra of the melanin-coated cotton was recorded to investigate the chemical modifications and compared with uncoated cotton as the control **(Figure 4C)**. Both samples exhibited similar characteristic peaks.^58^ In addition, the uncoated cotton displayed a peak at 1713 cm^-1^, which shifted towards a lower wavenumber (∼1600 cm□^1^) in the coated sample. This is attributed to the potential interaction of Al^3+^ ions with the carboxyl and phenolic functional groups present in tannic acid and melanin. The XPS spectra of uncoated cotton displayed characteristic peaks of cellulose with C 1s (285 eV) and O 1s (532 eV) (**Figure 4D and S6**).^59^ In contrast, the melanin-coated cotton indicated C 1s and O 1s along with additional peaks for N 1s (399 eV) and Na 1s (1072 eV) **(Figure 4D and S7)**.^60,61^ The appearance of the N 1s peak confirms the successful coating of melanin on the cotton, whereas the Na 1s signal is likely associated with residual reagents remaining from the post-treatment processes. Finally, the photothermal performance of melanin-coated cotton samples was evaluated using uncoated cotton as a control **(Figure 4E)**. It was observed that for the control samples under varying power densities (0.014 W/cm^-2^, 0.067 W/cm^-2^ and 0.5 W/cm^-2^) only minimal temperature rise was observed up to 32.5 °C. In contrast, the melanin-coated cotton samples exposed to the highest power density exhibited a temperature rise of up to 70.7 °C, indicative of the role of melanin deposited over the surface as an effective phothermal converter **(Figure 4F)**.

**Figure 4.**
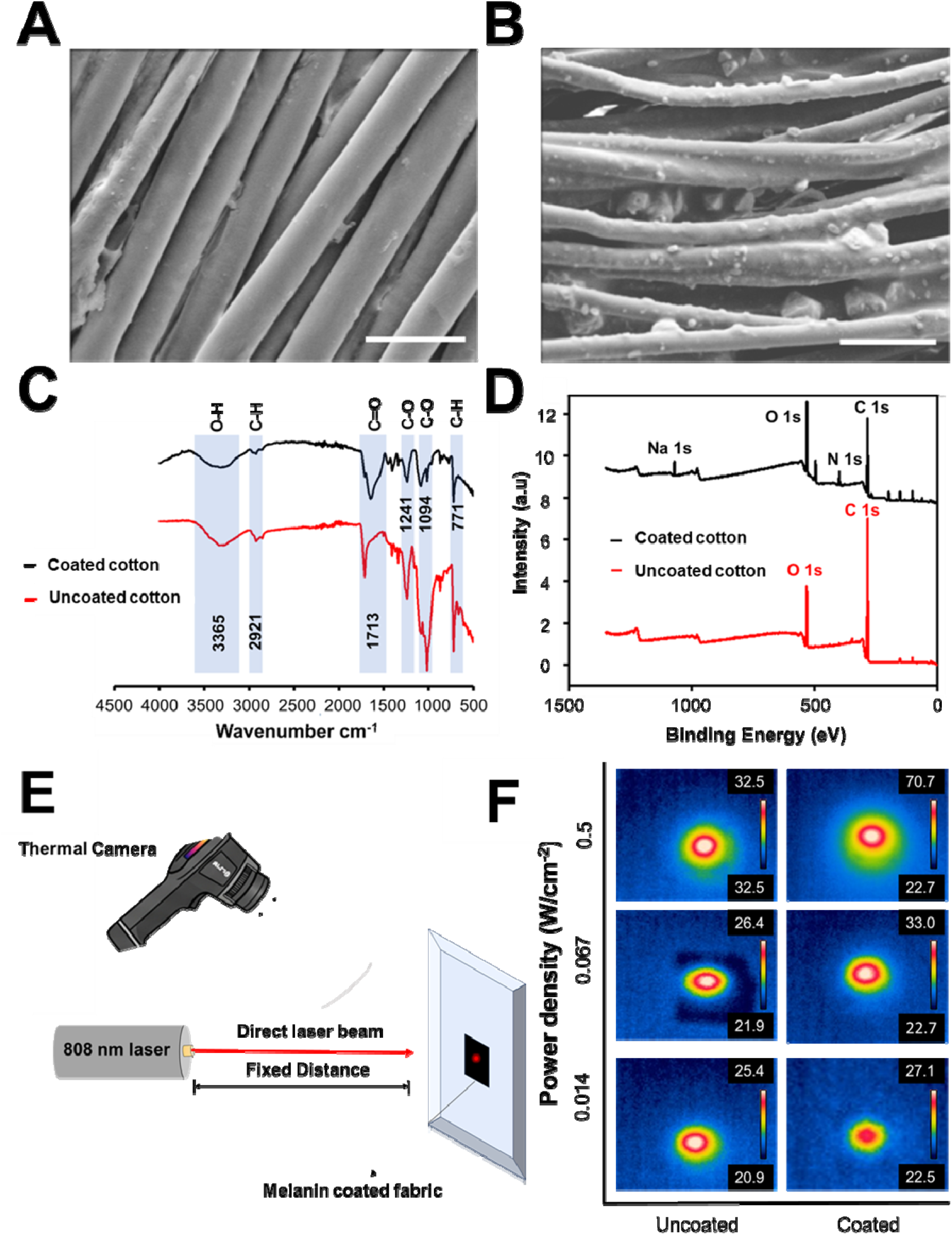
Characterization of melanin-coated cotton and photothermal activity of cotton. **(A)** SEM image of uncoated cotton showing well-organized cylindrical cellulose, **(B)** SEM image of melanin-coated cotton depicting non-homogeneous cellulose structures, **(C)** FTIR spectra of uncoated and coated samples showing characteristic cellulose peaks, with noticeable peak shift in the coated samples, **(D)** XPS spectra of uncoated and coated cotton showing C 1s and O 1s peaks in both samples, while the coated sample additionally exhibits N 1s and Na 1s signal, **(E)** Schematic illustration of the experimental setup used for photothermal evaluation of melanin-coated cotton under irradiation using an 808 nm near-infrared (NIR) laser for 6 min and its thermal imaging and **(F)** Infrared thermal images showing the temperature rise of melanin-coated cotton and the uncoated cotton at varying power densities (0.014 W/cm^-2^, 0.067 W/cm^-2^ and 0.5 W/cm^-2^). The scale bar in the SEM is 10 µm.

Following the successful coating of cotton, the developed methodology was further extended to wood to evaluate its applicability on alternative lignocellulosic materials. Wood, a lightweight, strong, and versatile biopolymeric material, is widely used in construction, packaging, and emerging functional applications.^62,63^ The wood samples were incubated in the melanin-containing growth media during IPTG induction, resulting in progressive black coating over the surface that intensified with increasing incubation time **(Figure 5A and 5B)**. We did not observe visible coating of the wood samples at the initial stages of melanin production (4 h and 8 h). After 12 h, a faint and patchy black coating became apparent, indicating the onset of melanin deposition. This coating intensified over 3 days of incubation. We explored two alternative approaches for coating wood, (1) where melanin production was allowed to occur in the *E*.*coli* culture (24 h), following which the wood was submerged and (2) the wood was incubated in the supernatant containing melanin. However, this method yielded a less intense coating than the *in situ* coating strategy **(Figure S8)**.

**Figure 5.**
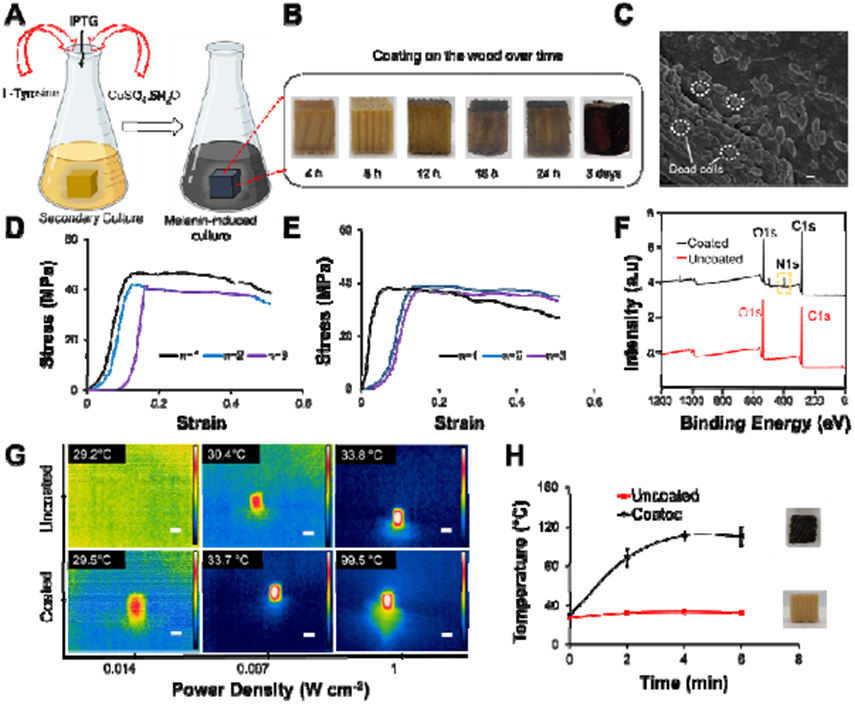
Methodology and characterization of melanin coated wood samples. **(A)** Schematic representation of preparing melanin coated wood using tyrosine (0.6 mM) and CuSO□·5H□O (30 µM) during IPTG induction, **(B)** Digital images of wood at different time intervals (4 h, 8 h, 12 h, 16 h, 24 h, and 3 days), showing progressive darkening due to melanin coating, **(C)** SEM image of wood after incubation in the culture media for 3 days indicating rod shaped *E*.*coli* cells. The scale bar used for the SEM images is 1 µm, **(D)** Stress–strain curves of LB media dipped wood under compression, **(E)** Stress–strain curves of melanin-coated wood under compression. n = 1,2 and 3 indicate the three independent experiments, **(F)** XPS spectra for coated and uncoated wood showing distinct signals for C1s and O1s with an additional N1s peak in coated wood suggesting the presence of melanin, **(G** Infrared thermographs showing temperature changes upon exposure to different laser power densities (0.014, 0.067, and 1 W cm□^2^) for uncoated wood (top row) and melanin coated wood (bottom row) and **(H)** Temperature vs. time graph comparing heating kinetics of uncoated (red) and melanin-coated (black) wood under laser exposure. The scale bar represents the length of the wood (1 cm).

Following melanin coating, the surface morphology of the coated wood was analyzed by SEM with an uncoated wood as the control. The melanin-coated wood revealed the presence of rod-shaped *E*.*coli* and uncoated wood indicated a smooth surface **(Figure 5C and S9)**.^64,65^ The mechanical strength was determined by a compression test, ensuring that the wood could withstand an adequate load. For the uncoated wood, the compressive strength evaluated was 50.69 ± 3.61 MPa and the Young’s modulus was 1.01 ± 0.18 GPa **(Figure S10)**. Similar tests were conducted on wood samples dipped in LB as the control and those coated with melanin **(Figure 5D and 5E)**. The compressive strength of LB dipped wood evaluated was 43.32 ± 2.90 MPa and the Young’s modulus was 0.84 ± 0.19 GPa. The compressive strength of melanin coated wood was 38.25 ± 0.75 and the Young’s modulus was 0.79 ± 0.24 GPa. XPS was conducted to determine the elemental composition of the wood surface. The XPS spectra for both the coated and the uncoated wood revealed characteristics signals corresponding to C 1s and O 1s **(Figure 5F)**. In addition, the coated wood showed an additional N 1s peak due to the nitrogen-containing moieties.^66^ The deconvolution of C 1s spectra for both uncoated and coated showed primarily three peaks, C_1_ peak corresponds to the C-H and C-C bond, C_2_ represents the O-C-O bond and C_3_ represents the O-C=O **(Figure S11A and S11B)**.^67^ Similarly, the deconvolution was done for O 1s for both showing primary peaks, O_1_ peak that corresponds to the C=O and O_2_ peak corresponds to the C-O bond **(Figure S11C and S11D)** The deconvolution of N 1s for the coated wood showed a peak for the presence of amine group **(Figure S12)**. FTIR spectra of the uncoated and coated wood samples were recorded to identify the functional groups involved in coating, while XRD analysis was performed to examine the crystalline nature of the melanin deposited on the surface **(Figure S13A and S13B)**.

The photothermal property of the wood was also determined using 808 nm laser with varying power densities (0.014, 0.067 and 1 Wcm^-2^). At the highest power density, 1 Wcm□^2^, for 6 min, the coated wood achieves a peak temperature of ∼100°C, compared to ∼34°C for the uncoated wood **(Figure 5G and S14)**. This substantial difference illustrates the ability of melanin coated wood to convert light to heat. Upon reducing the power density to 0.067 W cm□^2^, the melanin-coated wood attained a temperature of ∼33.5 °C demonstrating its effective photothermal performance even under lower irradiation intensity. At the lowest power density of 0.014 W cm□^2^, both samples (29.2°C for uncoated and 29.5°C for melanin coated) remain closer to ambient temperature. So, the time-dependent temperature profiles for coated and uncoated wood under a power density of 1 W cm^-2^ were analyzed to check the change in temperature with time **(Figure 5H)**. The coated wood rapidly heats up within the first 6 minutes, reaching a temperature plateau around 110 ± 9.57 °C, whereas the uncoated wood temperature shows a minimal increase. These results further highlight the coloration property of melanin and its potential for thermal management application.

## CONCLUSION

We have developed a whole-cell biocatalytic strategy for *in situ* synthesis and direct application of melanin for coating on material surfaces. The tyrosinase-mediated oxidation of L-tyrosine (substrate) in the presence of copper ions (co-factor) produces melanin that simultaneously coats the material surfaces submerged in the medium. This eliminated the need for pigment extraction and purification. The *E*.*coli* was inoculated until OD_600_ of 0.6-0.8, followed by the expression of tyrosinase in the presence of IPTG. The tyrosinase expression was optimized by varying the IPTG, L-tyrosine, and copper sulfate concentrations. For coating cotton samples, the pre-treatment process was systematically optimized by varying the presence of tannic acid, potash alum, and acetic acid. The results revealed that all three components were essential for achieving a uniform coating across the cotton surface. Furthermore, to improve the color fastness, the coated cotton samples were subjected to post-treatment comprising three cycles of coating, drying, and heat pressing. This resulted in improved resistance against water and detergent washing. The scanning electron micrographs of the uncoated cotton exhibited characteristic cylindrical fibril structures, whereas the coated samples displayed similar structures with an uneven surface likely due to melanin deposition across the surface. Subsequently, XPS analysis of the coated and uncoated samples revealed characteristic C 1s and O 1s peaks, while the coated cotton exhibited additional N 1s and Na 1s peaks confirming successful coating. The photothermal activity of the coated samples was investigated, showing a temperature increase up to 70 °C, whereas the uncoated samples exhibited only a minimal rise in temperature. This developed methodology was further extended to wood to assess its applicability to other lignocellulosic materials. Overall, the work establishes a simplified and potentially scalable route towards sustainable pigment production through living whole-cell system.

## AUTHOR CONTRIBUTIONS

K.P. conceived the original idea and planned the experiments. A.D. and T.T. carried out experiments and data analysis. K.P. supervised the research along with providing feedback during the writing of the manuscript. All authors provided critical feedback and helped shape the research, analysis, and manuscript.

## ACKNOWLEDGEMENTS

The authors thank CRTDH (Common Resource and Technology Development Hub) (Professor Chinmay Ghoroi), Nanoplasmonic Research Lab (Professor Saumyakanti Khatua) and CIF (Central Instrumentation Facility) for providing various instrumentation facilities at IIT Gandhinagar. The authors would also like to thank the carpentry workshop for providing equipment for sample preparation.

## Table of Content

Whole-cell biocatalytic melanin synthesis using recombinant E. coli for direct, in situ coating of cotton and enhanced photothermal activity.

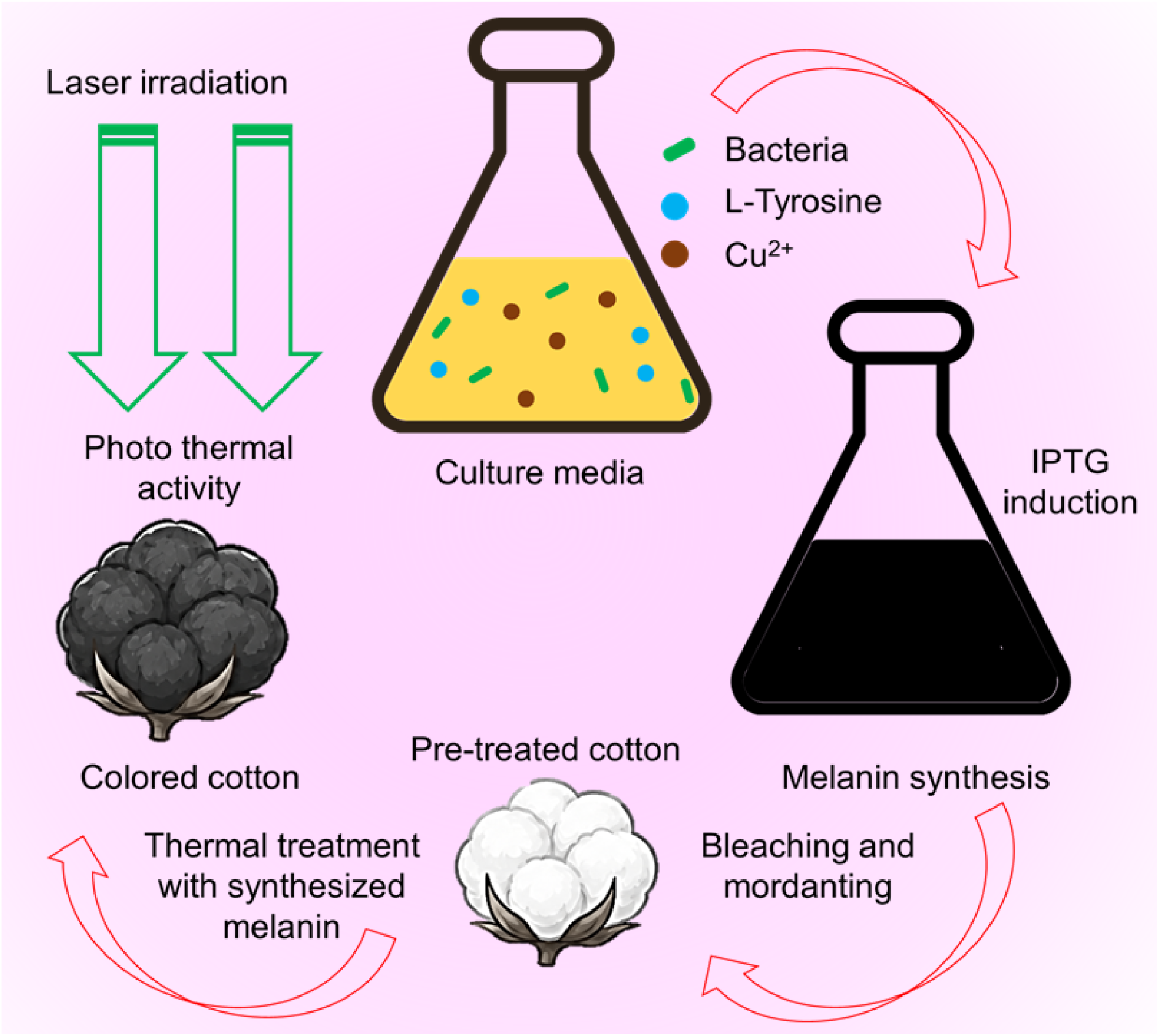

