## Supplementary material for "In Situ Biocatalytic Color Generation Using Living Whole-Cell Systems for Sustainable Textile Dyeing": SI

Karthik Pushpavanam, Ph.D.

Chemical Engineering

IIT Gandhinagar

Gujarat, India, 382355

**Table S1. Nucleotide sequence and amino acid sequence of pLN-AIDA-Tyr1.**

| **>Nucleotide sequence** |
|---|
| atgaataaggcctacagtatcatttggagccactccagacaggcctggattgtggcctcagagttagccagaggacatggttttgtccttg<br>caaaaaatacactgctggtattggcggttgtttccacaatcggaaatgcatttgcagtcgaccaccatcaccatcaccatctggaagcgct<br>gttccagggtccgggtaccggtaataaatatcgcgttcgtaaaaatgtgttgcatctgaccgacaccgaaaaacgcgatttcgtgcgtacc<br>gttctgattctcaaagagaaaggtatctacgaccgctatatcgcctggcatggcgcagcgggtaaatttcatactcctcctggcagcgatc<br>gcaacgccgcgcatatgagcagcgccttcctgccgtggcaccgcgaatatctgctgcgttttgaacgcgatctgcaaagcatcaaccca<br>gaagtgaccctgccgtactgggagtgggaaaccgatgcgcaaatgcaggatccgagtcagtcgcagatctggtcggcggactttatgg<br>gcggcaacggtaacccgatcaaagattttattgtcgataccggtccgttcgccgcaggccgctggaccaccattgatgaacagggcaat<br>ccgagcggcggcctgaagcgcaattttggcgccaccaaagaagcgccgacgctgccgacccgagatgacgtgctgaacgcgctgaa<br>aattacccagtatgacacgccgccgtgggatatgaccagccagaatagctttcgtaaccaactggaaggttttatcaatggcccgcagct<br>gcacaaccgcgtgcatcgttgggttggtggtcaaatgggcgtagtcccaactgcgccgaacgatccagtatttttcctgcatcacgccaa<br>cgtggatcgaatctgggcggtctggcagattattcacaggaatcagaattatcagccaatgaaaaacggtccgttcggtcagaacttccg<br>cgacccgatgtatccgtggaacaccaccccagaagatgttatgaaccatcgcaaactgggctatgtctacgatatcgaactgcgcaaaa<br>gcaaacgttccag |
| **>Amino acid sequence** |
| MNKAYSIIWSHSRQAWIVASELARGHGFVLAKNTLLVLAVVSTIGNAFAVDHHHHHH<br>LEALFQGPGTGNKYRVRKNVLHLTDTEKRDFVRTVLILKEKGIYDRYIAWHGAAGKF<br>HTPPGSDRNAAHMSSAFLPWHREYLLRFERDLQSINPEVTLPYWEWETDAQMQDPSQ<br>SQIWSADFMGGNGNPIKDFIVDTGPFAAGRWTTIDEQGNPSGGLKRNFGATKEAPTLP<br>TRDDVLNALKITQYDTPPWDMTSQNSFRNQLEGFINGPQLHNRVHRWVGGQMGVVP<br>TAPNDPVFFLHHANVDRIWAVWQIIHRNQNYQPMKNGPFGQNFRDPMYPWNTTPED<br>VMNHRKLGYVYDIELRKSKRS |

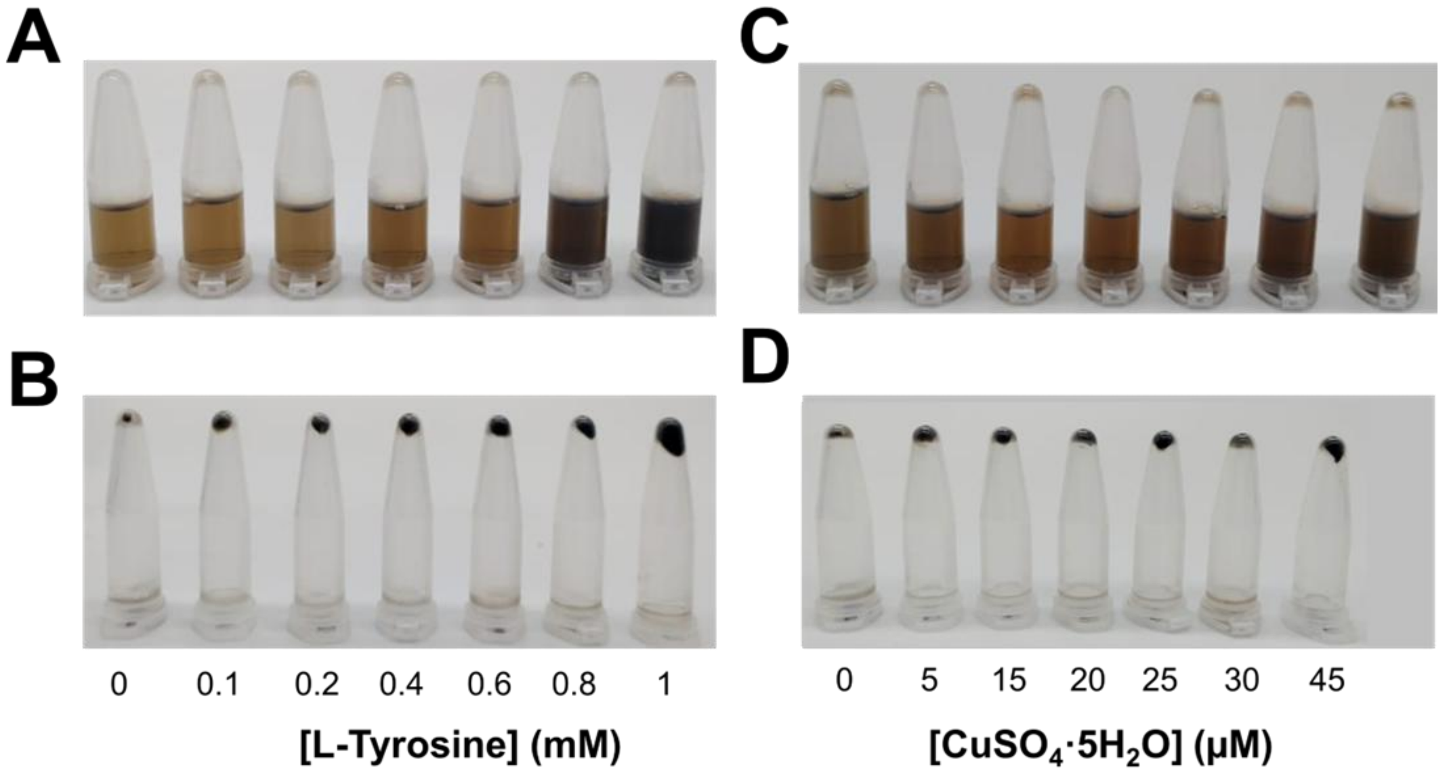


**Figure S1. Effect of tyrosine and $CuSO_4.5H_2O$ on melanin production. (A)** The supernatant of the growth media with increasing concentration of tyrosine (0 to 1 mM) and fixed concentration of IPTG (1 mM) and $CuSO_4.5H_2O$ (30 µM) indicating an increase in the intensity of black color, **(B)** The corresponding cell pellets with increasing concentration of tyrosine showing an increase in black coloration, **(C)** The supernatant of the growth media with increasing concentration of $CuSO_4.5H_2O$ (0 to 45 µM) and fixed concentration of IPTG (1 mM) and tyrosine (0.6 mM) displays indicating an increase in the intensity of black color and **(D)** The corresponding cell pellets indicating with increasing $CuSO_4.5H_2O$ concentration showing an increase of black coloration.

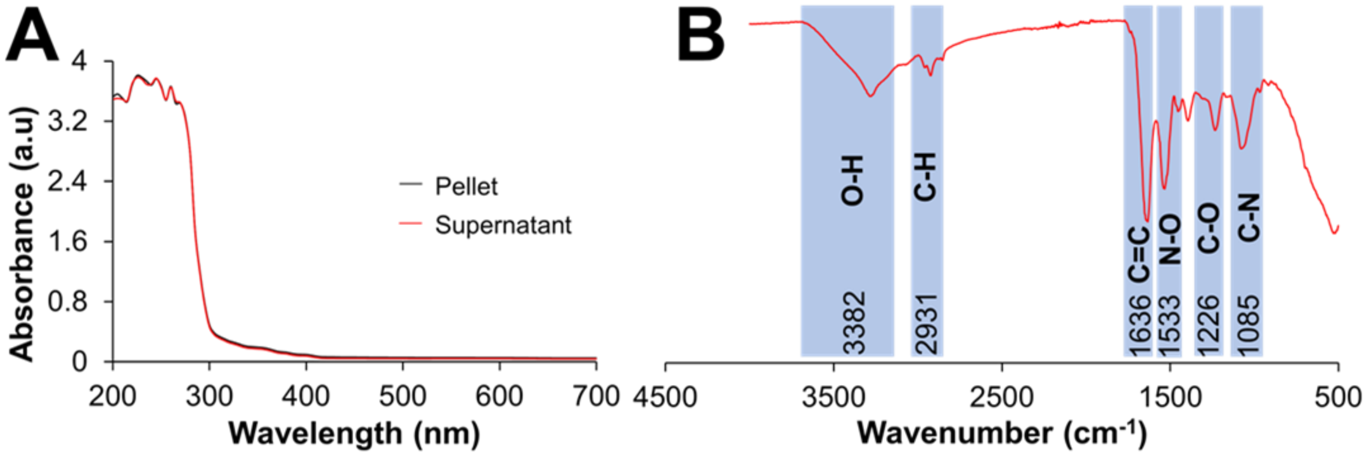


**Figure S2. Characterization of synthesized melanin. (A)** UV–visible absorption spectra (200–700 nm) of the isolated melanin pellet and the corresponding supernatant, and **(B)** FTIR spectra of the synthesized melanin showing characteristic absorption bands corresponding to the functional groups present within melanin.

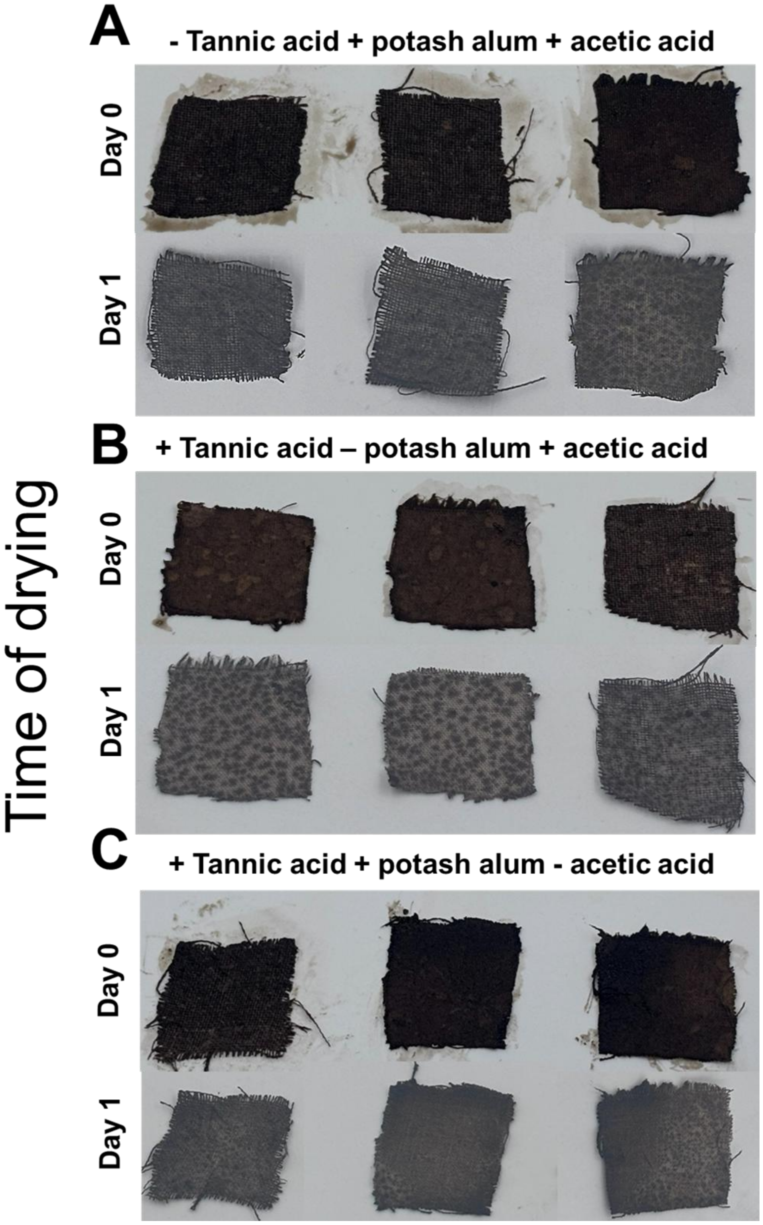


**Figure S3. The effects of tannic acid, potash alum, and acetic acid pretreatment on the uniformity and darkness of melanin coated cotton samples.** Pictures showing the representative appearance of melanin-coated cotton before drying (Day 0) and after drying (Day 1) from three independent experiments. **(A)** In the absence of tannic acid, but in the presence of potash alum and acetic acid, an intense black coloration was initially observed without drying but after drying the sample exhibited a light grey, patchy appearance, **(B)** In the absence of potash alum but in the presence of tannic acid and acetic acid, an intense black coloration was initially observed without drying but after drying exhibited a similar light grey, patchy appearance and **(C)** In the absence of acetic acid but in the presence of tannic acid and potash alum, an intense black coloration was initially observed without drying and after drying a light grey, patchy appearance was observed.

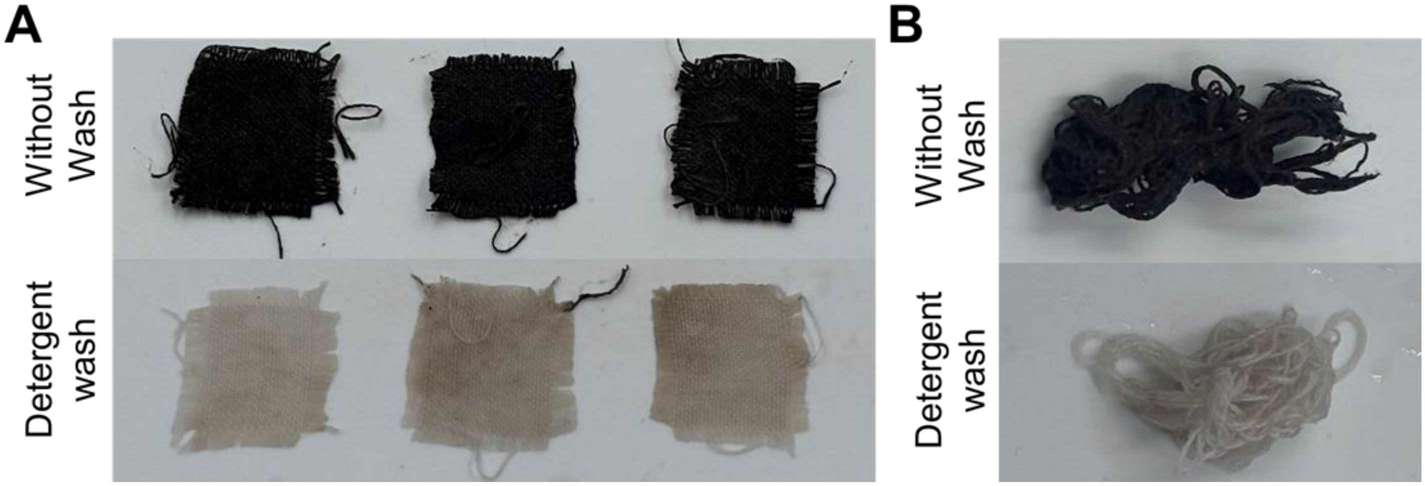


**Figure S4. The color fastness in melanin-coated cotton cloth and cotton thread in the absence of post treatment process.** These are representative pictures of three independent experiments. **(A)** Cotton fabric samples coated with melanin without any post-treatment exhibited intense black coloration; however, complete loss of color was observed after a single detergent washing cycle and **(B)** Similarly, the cotton thread coated with melanin without any post-treatment exhibited intense black coloration; however, complete loss of color was observed after a single detergent washing cycle.

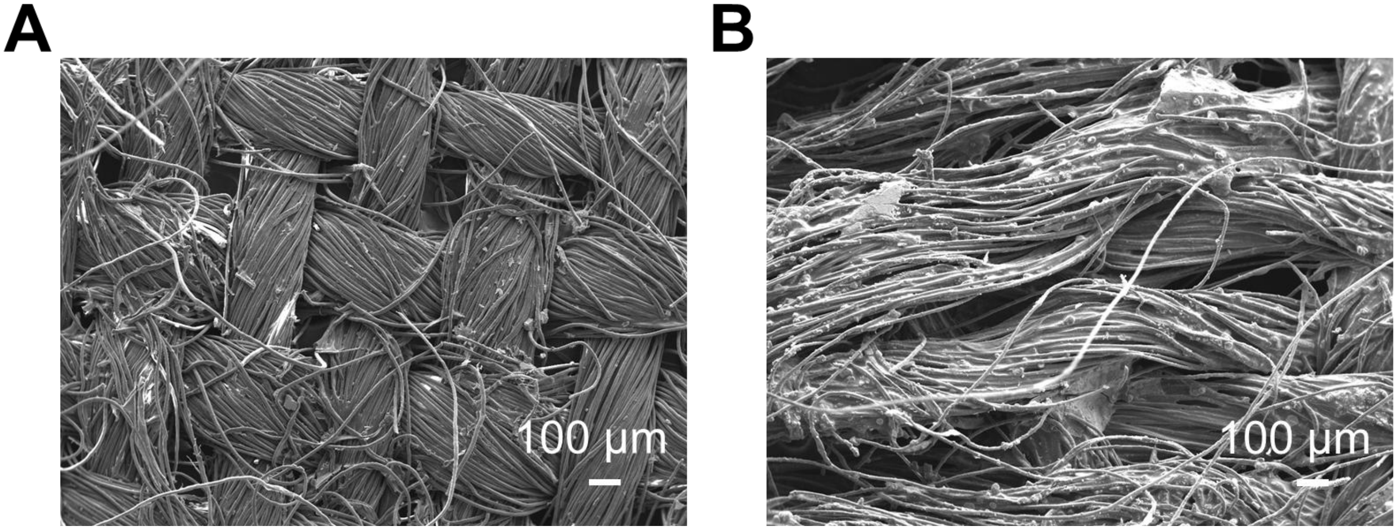


**Figure S5. Scanning electron microscopic images of uncoated and melanin-coated cotton samples.** **(A)** SEM image of uncoated cotton showing arranged cylindrical structures of fibers, and **(B)** SEM image of the melanin–coated cotton showing partially loosened cylindrical structures of fibers with uneven particulate deposits likely due to melanin deposition.

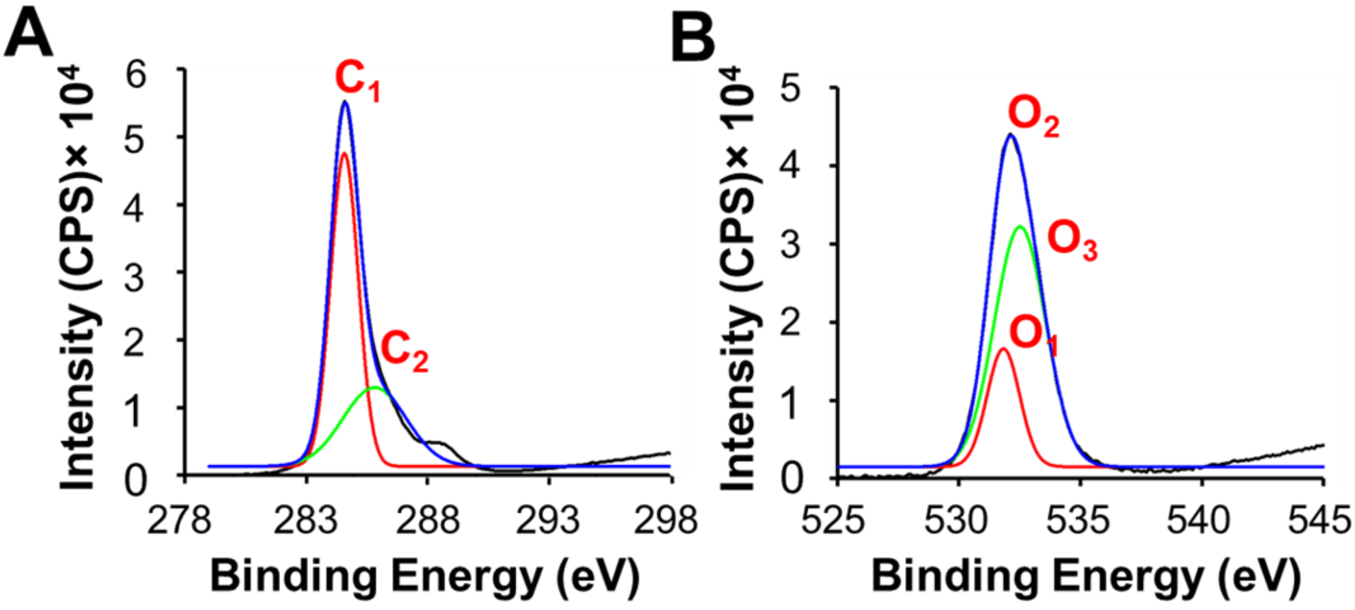


**Figure S6. XPS spectra of C 1s and O 1s regions for the uncoated cotton sample**. **(A)** Deconvoluted C 1s spectra with peaks labeled as $C_1$ for C-H and $C_2$ for O-C-O bonds and **(B)** Deconvoluted O 1s spectra with peaks labeled as $O_1$ and $O_2$ corresponds to C=O and $O_3$ corresponds to C-O.

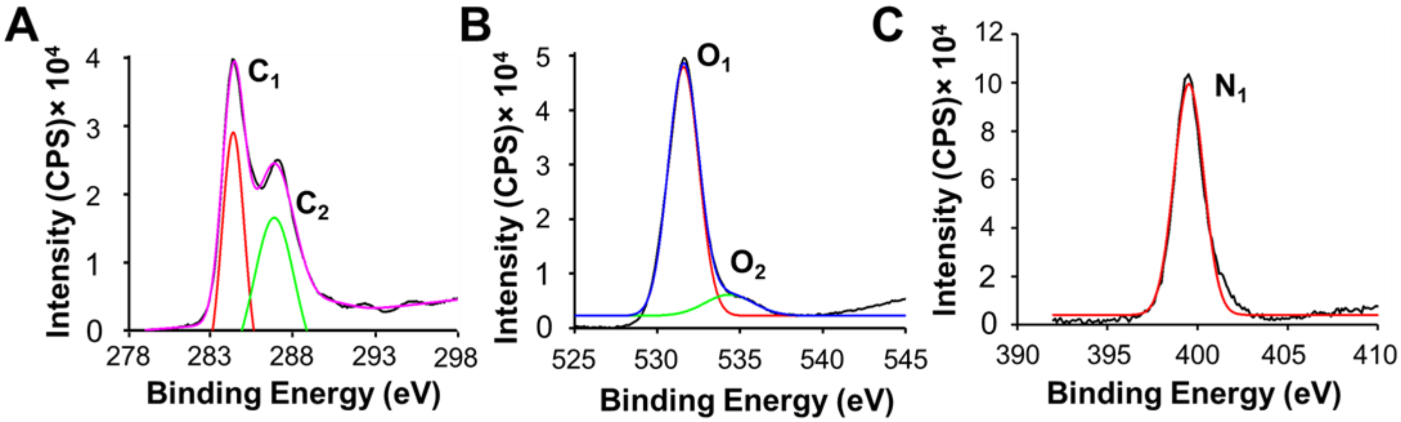


**Figure S7. XPS spectra of C 1s, O 1s and N 1s regions for melanin coated cotton samples**. **(A)** Deconvoluted C 1s spectra with peaks labeled as $C_1$ for C-H and $C_2$ for O-C-O bonds, **(B)** Deconvoluted O 1s spectra with peaks labeled as $O_1$ and $O_2$ corresponds to C=O bonds and **(C)** Deconvoluted N 1s spectra with a single fitted peak indicating $–NH_2$ bonds due to the melanin deposition.

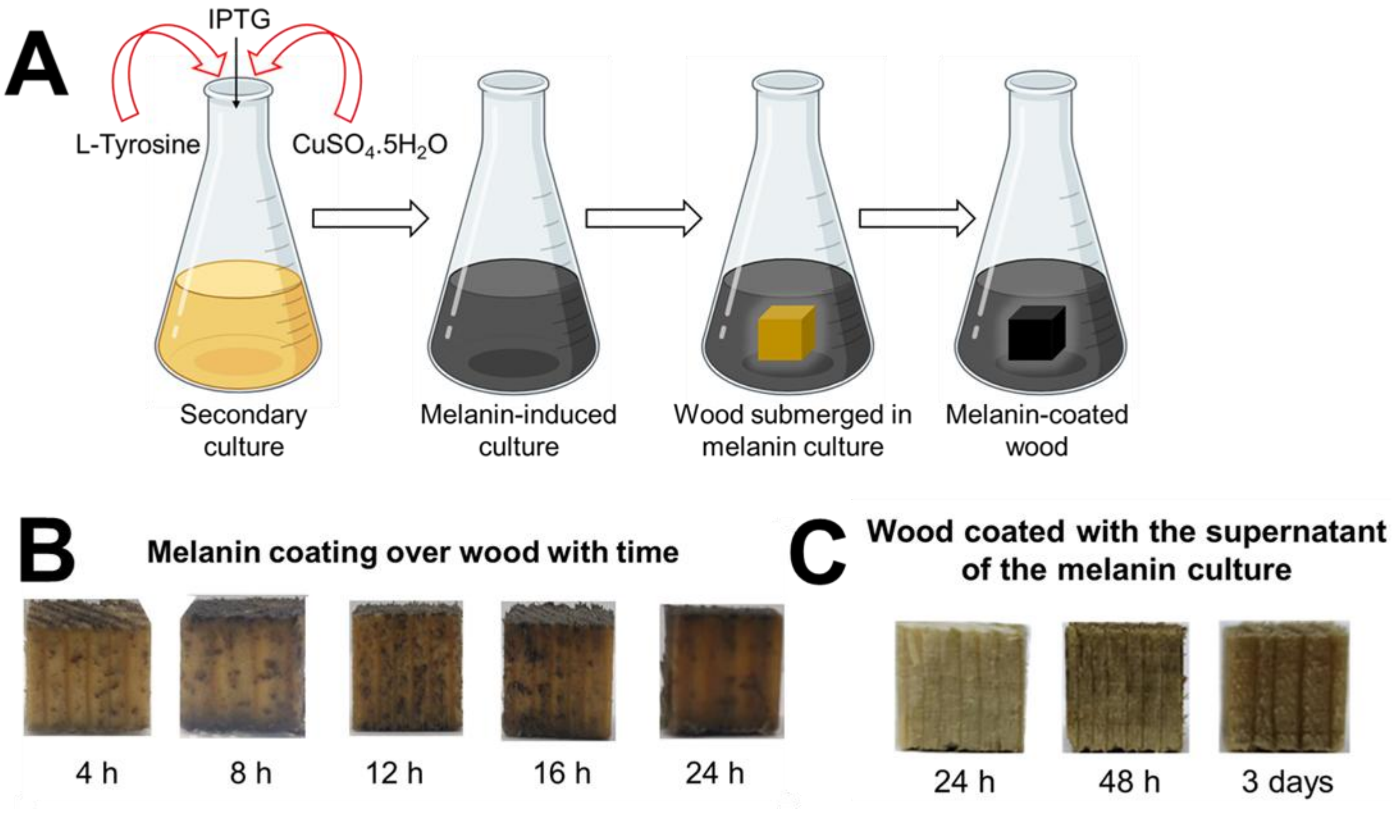


**Figure S8. An alternative methodology for melanin coating on wood**. **(A)** Schematic representation of the methodology for coating wood with IPTG-inducible melanin in the presence of tyrosine and $CuSO_4 \cdot 5H_2O$, where the wood samples were submerged in the culture after 24 h of IPTG induction and **(B)** The visual progression of melanin coating over time over the wood surface resulting in black patchy coating after 24 h and **(C)** Digital images of wood coated with the supernatant of the melanin culture, resulting in a less intense black coloration.

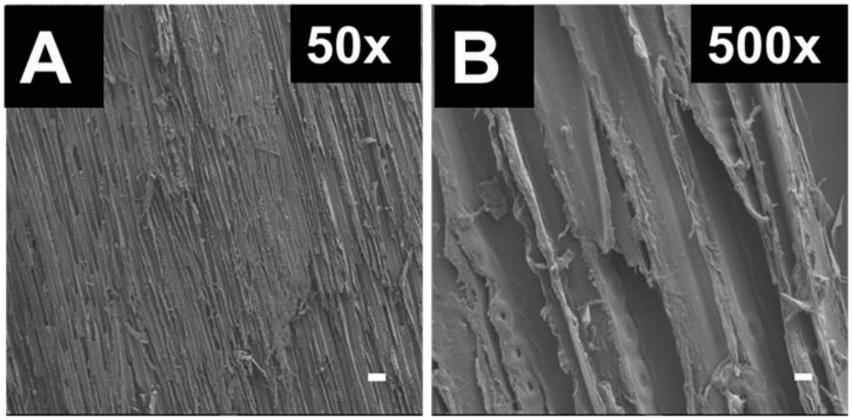


**Figure S9. Scanning electron micrographs of and coating done by the supernatant of the melanin culture. (A)** SEM images of raw wood at 50x magnification indicating rough fibrous surface, **(B**) SEM images of the raw wood at 500x with roughness over the surface.

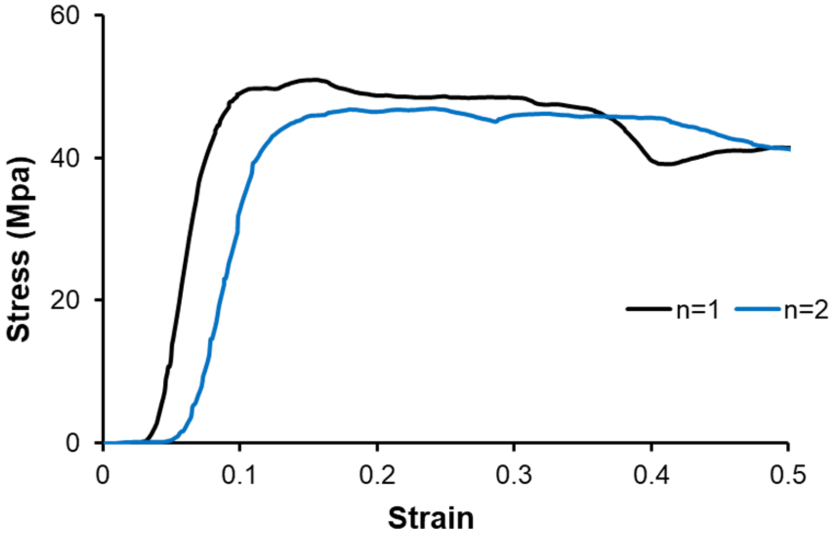


**Figure S10.** Stress–strain curves under compression of uncoated wood submerged in water evaluated for two independent experimental trials.

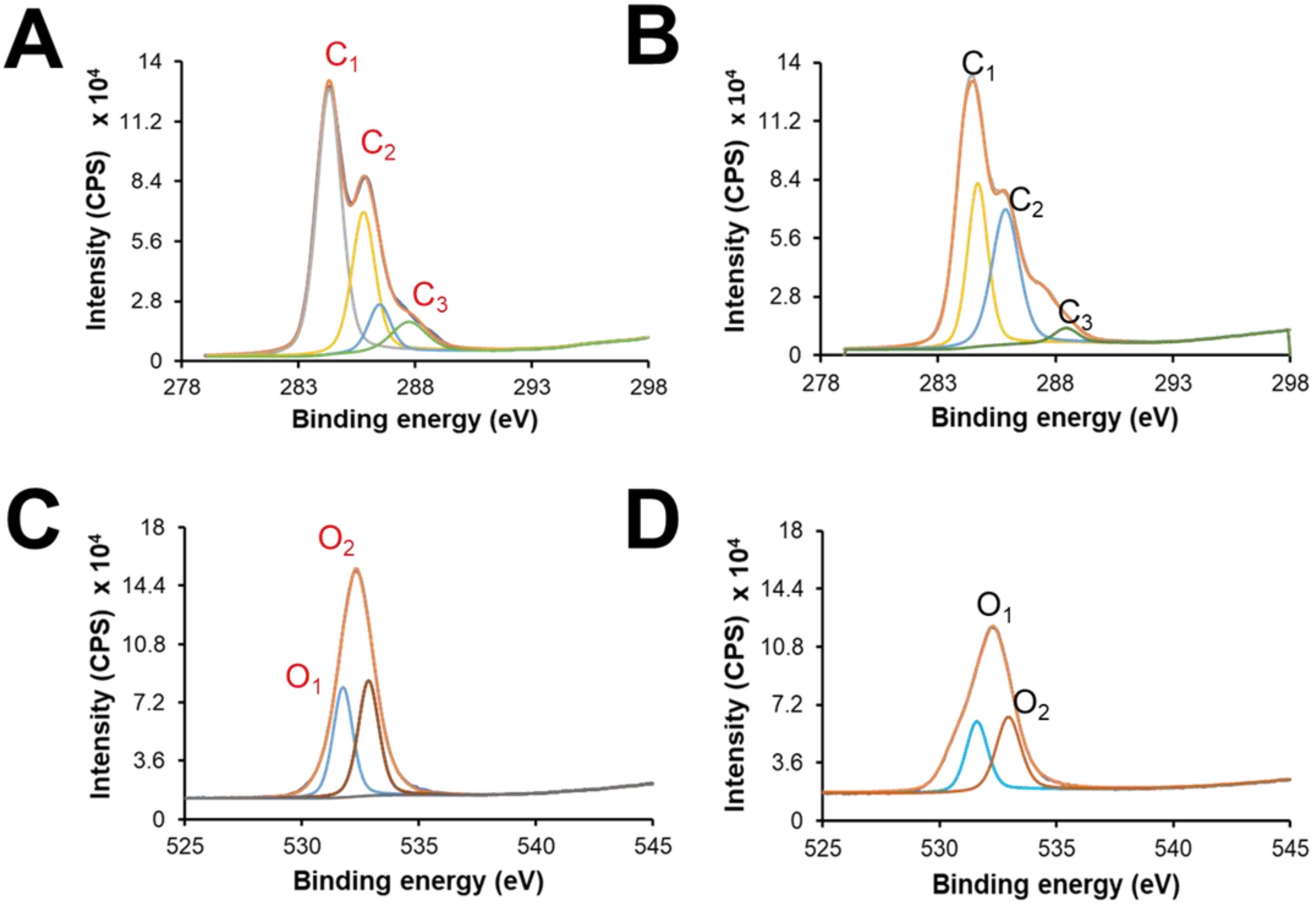


**Figure S11. XPS spectra of C 1s and O 1s regions for wood surfaces before and after melanin coating**. **(A)** Deconvoluted C 1s spectra of uncoated wood with peaks labeled as $C_1$ and $C_2$ corresponding to C-H and O-C-O bonds and $C_3$ corresponding to O-C=O bond, **(B)** Deconvoluted C 1s spectra of coated wood with peaks labeled as $C_1$ and $C_2$ corresponding to C-H and O-C-O bonds respectively and $C_3$ corresponding to O-C=O bond, **(C)** Deconvoluted O 1s spectra of uncoated wood with peaks labeled as $O_1$ corresponding to C=O, and $O_2$ corresponds to C=O bond and **(D)** Deconvoluted O 1s spectra of coated wood with peaks labeled as $O_1$ corresponding to C=O, and $O_2$ corresponds to C=O bond.

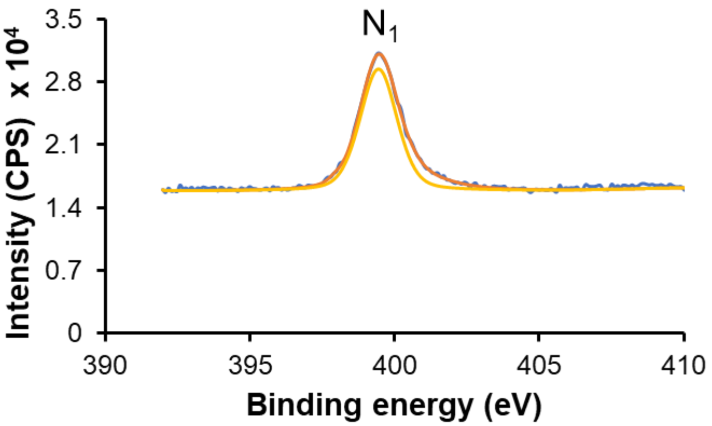


**Figure S12. Deconvoluted N 1s peak of the XPS spectra of melanin-coated wood.** It shows a single fitted peak, indicating successful surface modification of the wood.

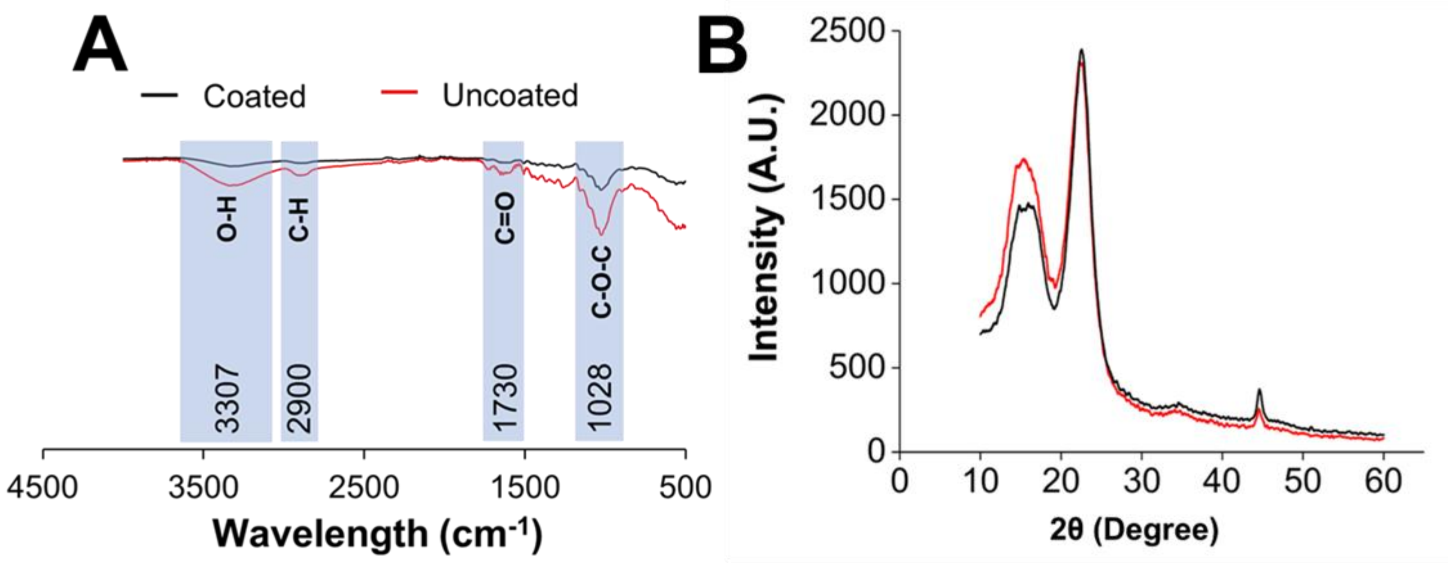


**Figure S13. Characterization of melanin coated and uncoated wood surfaces. (A)** FTIR spectra between 4000 to 500 cm$^{-1}$ for both the coated and uncoated wood indicated characteristics peaks of wood at 3307 cm$^{-1}$, 2900 cm$^{-1}$, 1730 cm$^{-1}$ and 1028cm$^{-1}$ and **(B)** XRD patterns of uncoated and coated wood indicating retention of the crystalline cellulose structure suggesting minimal changes to wood crystallinity.

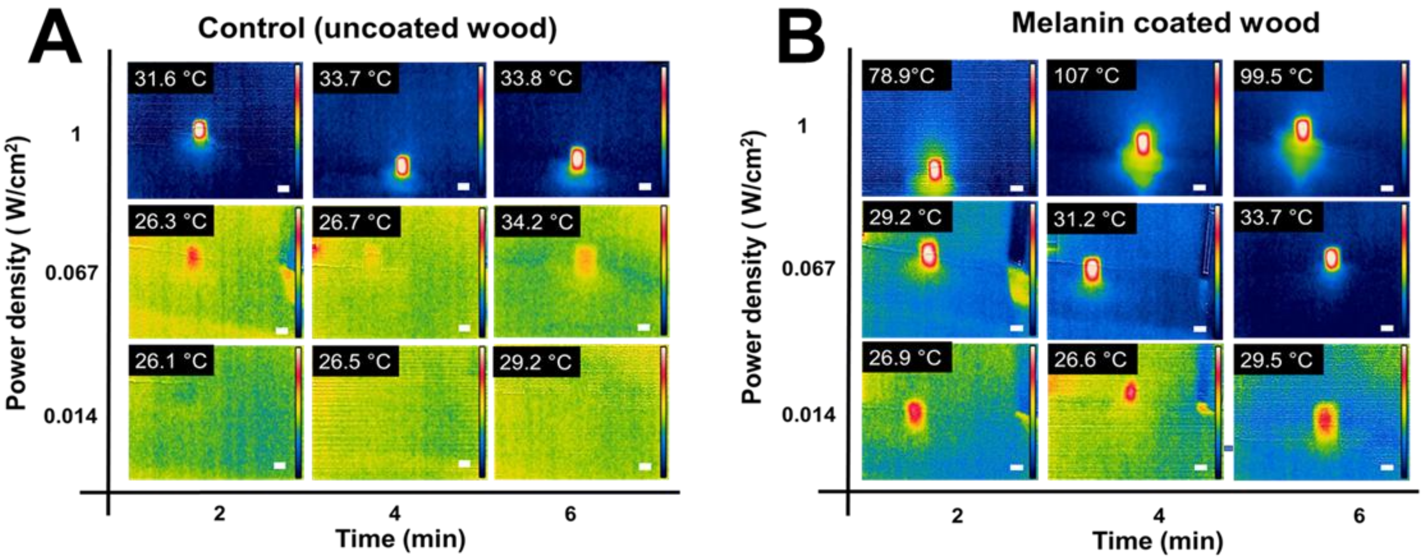


**Figure S14. Photothermal activity of the melanin-coated and uncoated wood at different laser irradiations. (A)** Infrared images of uncoated wood showing temperature changes upon exposure to 808 nm laser with different power densities (0.014, 0.067, and 1 W $cm^{-2}$) for time intervals, 2,4, and 6 min indicating minimal temperature increase and **(B)** Infrared images of coated wood showing temperature changes upon exposure to an 808 nm laser at different power densities (0.014, 0.067, and 1 W $cm^{-2}$) for 2, 4, and 6 min, indicating increased temperature with increasing power density. The scale bar in the images represents the length of the wood (1 cm).
